# Patient-derived organoids identify tailored therapeutic options and determinants of plasticity in sarcomatoid urothelial bladder cancer

**DOI:** 10.1101/2023.02.20.528313

**Authors:** Michele Garioni, Lauriane Blukacz, Sandro Nuciforo, Romuald Parmentier, Luca Roma, Mairene Coto-Llerena, Heike Pueschel, Salvatore Piscuoglio, Tatjana Vlajnic, Frank Stenner, Hans-Helge Seifert, Cyrill A. Rentsch, Lukas Bubendorf, Clémentine Le Magnen

## Abstract

Sarcomatoid Urothelial Bladder Cancer (SARC) is a rare and aggressive histological subtype of bladder cancer for which therapeutic options are limited and experimental models are lacking. Here, we report the establishment of the first long-term 3D organoid-like model derived from a SARC patient (*SarBC-01*). SarBC-01 emulates aggressive morphological and phenotypical features of SARC and harbor somatic mutations in genes frequently altered in sarcomatoid tumors such as *TP53, RB1*, and *KRAS*. High-throughput drug screening, using a library comprising 1567 compounds in SarBC-01 and organoids derived from a patient with conventional urothelial carcinoma (UroCa), identified drug candidates active against SARC cells exclusively, or UroCa cells exclusively, or both. Among those, standard-of-care chemotherapeutic drugs inhibited both SARC and UroCa cells, while a subset of targeted drugs was specifically effective in SARC cells, such as agents targeting the Glucocorticoid Receptor (GR) pathway. In two independent patient cohorts, GR was found to be significantly more expressed, at mRNA and protein level, in SARC as compared to UroCa tumor samples. Further, glucocorticoid treatment impaired the mesenchymal morphology, abrogated the invasive ability of SARC cells, and led to transcriptomic changes associated with reversion of epithelial-to-mesenchymal transition, at single-cell level. Altogether, our study highlights the power of organoids for precision oncology and for providing key insights into factors driving rare tumor entities.

## Introduction

While more than 90% of bladder cancers present as “conventional” urothelial carcinomas (UroCa), additional histological subtypes have been described which are often clinically aggressive and challenging to treat (1). Among those subtypes, sarcomatoid urothelial bladder cancer (SARC) typically associates with early metastatic spread to distant organs and is linked to poor prognosis. Given that SARC is often found concomitant to conventional UroCa, it has been proposed that both components may share a common ancestor. Supporting this hypothesis, recent sequencing efforts have suggested that SARC exhibit genomic features common with UroCa and may evolve via the dysregulation of epithelial-mesenchymal transition (EMT) pathways (2,3). Yet SARC tumors differ from conventional UroCa by an enrichment of *TP53, RB1* and *PIK3CA* mutations and an enhanced expression of genes linked to chromatin remodelling and EMT (2). In light of these specific features, a consensus for the correct clinical management of this rare histological subtype is still lacking.

The recent advent of advanced *in vitro* models such as patient-derived organoids (PDOs) has opened new avenues to decipher the pathogenesis and therapeutic vulnerabilities of a wide range of tumors, including bladder cancer (4,5). A lack of experimental models emulating rare entities, such as SARC, however hampers the efforts to decipher mechanisms driving such diseases and the development of tailored clinical strategies. Here, we established a long-term *in vitro* PDO model from a patient harboring SARC (***SarBC-01***), characterized its phenotypical features *in vitro* and *in vivo*, and defined its genomic and drug sensitivity profile (workflow summarized in **Fig. 1**). Based on the drug sensitivity profile, we identify the glucocorticoid receptor (GR) to be highly expressed in two cohorts of SARC tumors and showed that treatment with glucocorticoids lead to phenotypic and transcriptomic changes associated with reversion of epithelial-to-mesenchymal transition.

**Figure 1.**
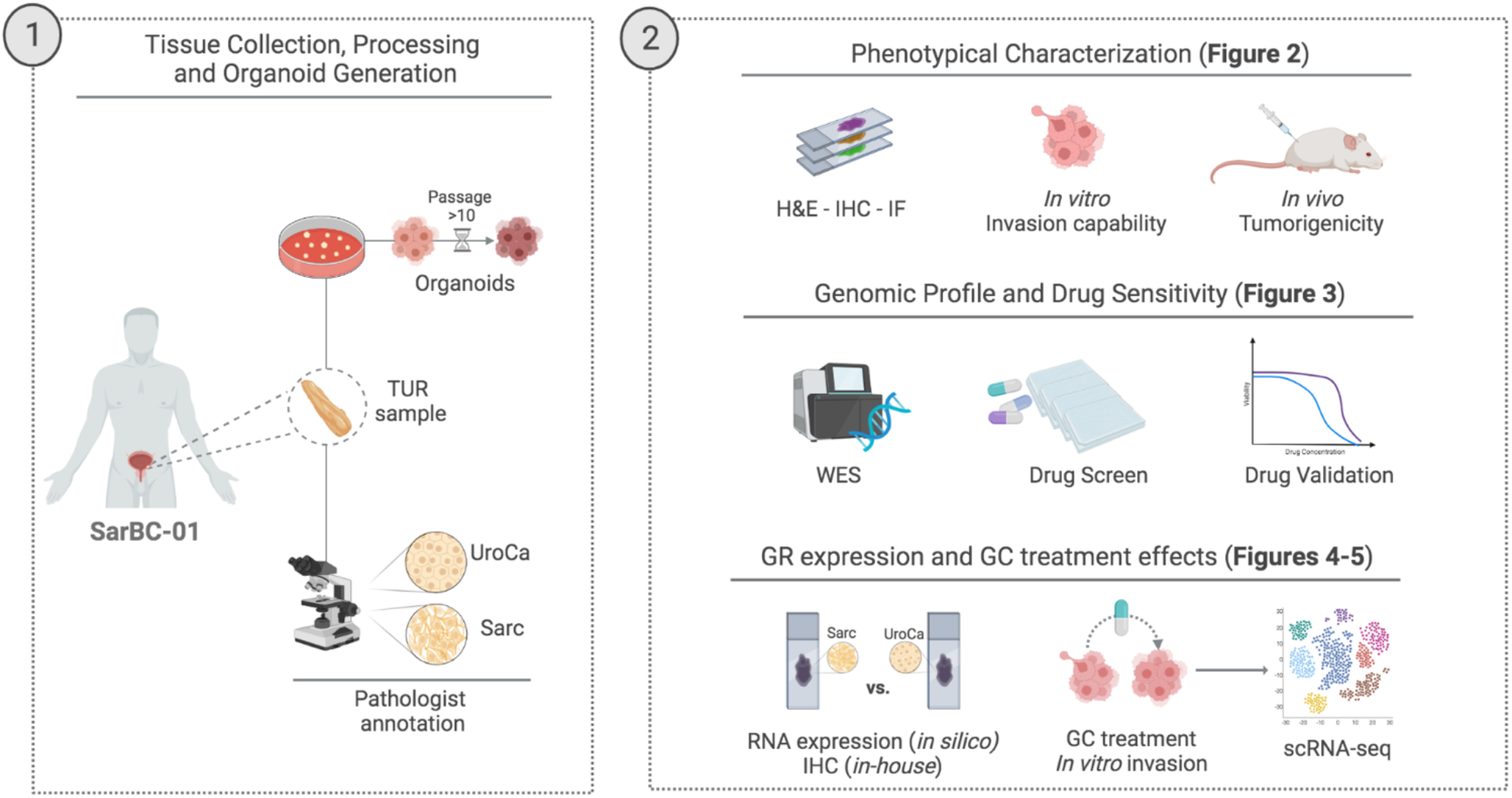
Cartoon depicting the distinct morphological components of the patients’ tumor and the experimental workflow used to generate organoids (1) and subsequent analyses (2). *UroCa: urothelial carcinoma; Sarc: sarcomatoid urothelial bladder cancer; TUR: transurethral resection; H&E: hematoxylin and eosin; IHC: immunohistochemistry; IF: immunofluorescence; WES: whole exome sequencing; GR: glucocorticoid receptor; GC: glucocorticoids*. Created with BioRender.com

## Results

### Generation of a long-term organoid model derived from a SARC patient, which retain phenotypic characteristics of SARC

We generated organoid cultures out of a fragment of a tumor derived from a SARC patient (***SarBC-01***), adapting a previously-published protocol (4). SarBC-01 organoids, were maintained in culture at long-term (>42 passages and >700 days in culture) and underwent multiple cycles of freezing and thawing, without notable difference of growth and morphology in culture (**Fig. 2A**). The original patient tumor displayed a wide spectrum of differentiation patterns ranging from poorly differentiated UroCa to SARC components, the latter being characterized by malignant cells with a spindle-like morphology (**Fig. 2B**). The UroCa component retained expression of epithelial markers such as E-cadherin and Cytokeratin (CK) 7, and expressed high levels of CD138 and GATA3. In contrast, these proteins were absent or expressed at very low level in the SARC counterpart, while CD44 and P53 were highly expressed in both components. Notably, “early” and “late” passage organoids (passage 6 and 20, respectively) displayed marker expression patterns consistent with a SARC-like phenotype, suggesting that they may derive from the SARC component or that this tumor component may have preferentially grown in culture (**Fig. 2B**). Similar to the SARC patients’ tumor component and in contrast to the UroCa component, organoids were negative for additional epithelial markers (CK5, CK8), yet showed a strong expression of Vimentin, hallmark of mesenchymal-like cells (**Fig. 2C**, left panel).

**Figure 2.**
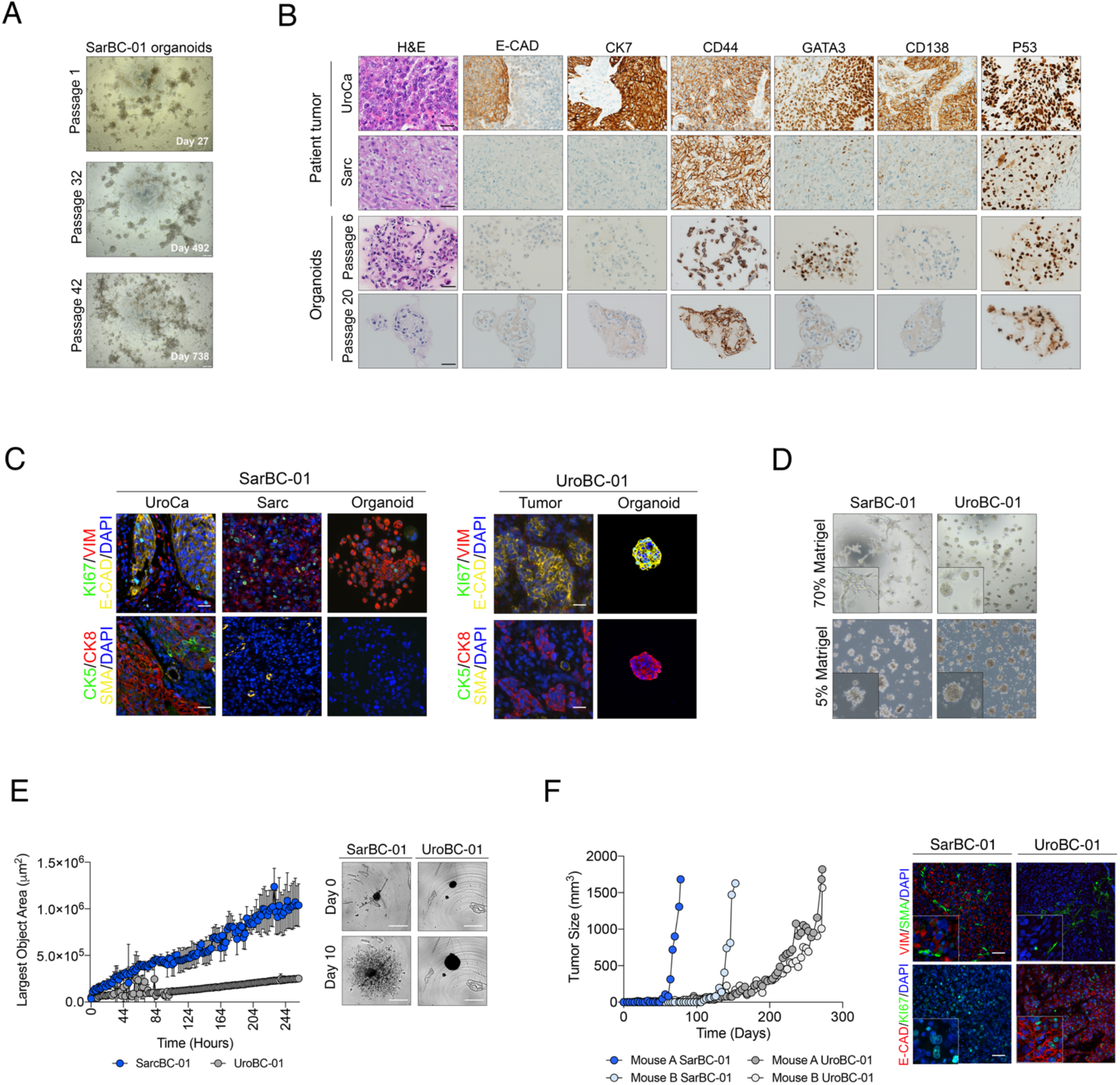
Generation of a long-term *in vitro* model derived from a SARC patient, which displays mesenchymal, invasive, and tumorigenic properties. **(A)** Representative bright field images of SarBC-01 organoids at passage 1 (27 days of culture), passage 32 (492 days of culture), and passage 42 (728 days of culture). **(B)** Phenotypic analyses of the distinct tumor components (UroCa, Sarc) and derived organoids at early and late passage (passage 6 and 20, respectively). Shown are representative H&E images and IHC staining for the indicated antibodies. Scale bars represent 50 μm **(C)** Immunofluorescence analyses of SarBC-01 (left) and UroBC-01 (right) tumor and organoids pairs. Shown are representative images for the indicated antibodies. *Vim: Vimentin, E-cad: E-cadherin, SMA: Smooth Muscle Actin*. DAPI: 4’,6-diamidino-2-phenylindole. Scale bars represent 50 μm **(D)** Representative bright-field images of organoids derived from SarBC-01 and UroBC-01 in presence of high (70%) or low (5%) Matrigel concentration. Insets show higher magnification of a representative region. **(E)** Left panel: Largest bright field object area measured over time for spheroids generated from SarBC-01 cells and UroBC-01 cells. Analyses were performed using a mixed-effects analysis (p=0.0001). Data are represented as means of three spheroids and the error bars represent standard deviations (SD). Two independent biological replicates were performed and one representative replicate is shown. Right panel: representative images of SarBC-01 and UroBC-01 spheroids at day 0 and day 10 are shown. Scale bars represent 800 μm **(F)** Left panel: Growth kinetic of xenografts generated by subcutaneous injection of SarcBC-01 and UroBC-01 cells in NSG mice. Two mice were injected per organoid line. Right panel: Immunofluorescence analyses of SarBC-01 (left) and UroBC-01 (right) derived xenografts. Shown are representative images for the indicated antibodies. *Vim: Vimentin, E-cad: E-cadherin, SMA: Smooth Muscle Actin*. Insets show higher magnification of a representative region. Scale bars represent 50 μm

### SarBC-01 organoids display aggressive mesenchymal features

To provide a comparative model of UroCa for further analyses, we used a novel long-term organoid line derived from a patient with conventional UroCa and generated in our laboratory (***UroBC-01***). In stark contrast with SarBC-01 cells, UroBC-01 organoids expressed high levels of epithelial markers (E-adherin, CK8) and lacked expression of Vimentin, consistent with an UroCa phenotype (**Fig. 2C**, right panel). Bright-field images of the two models highlighted the invasive potential of SarBC-01, which formed networks of spindle-like cells in the presence of Matrigel, in contrast to UroBC-01 which retained its spherical morphology (**Fig. 2D**). Consistent with these observations, SarBC-01 displayed a significantly higher invasive capacity *in vitro*, as compared to UroBC-01 (mixed-effects analysis, p=0.0001; **Fig. 2E**). These *in vitro* invasive features correlated with a faster tumorigenic capacity *in vivo* in the SarBC-01 model (2/2 xenografts generated for each model, average time between palpable tumor and end-point of 35 days in SarBC-01 vs. 210 days in UroBC-01; **Fig. 2F**). Xenografts generated out of the SarBC-01 displayed rhabdoid features and high expression of Vimentin, while UroBC-01-derived xenografts were negative for this marker (cells with human origin; **Fig. 2F** and **Supplementary Fig. 1**). Notably, portions of xenografts derived from SarBC-01 cells displayed intermediate phenotypes with poorly differentiated UroCa-like morphology and partial recovery of expression of epithelial markers, reflecting the spectrum of heterogeneity found in the patient sample and suggesting that SarBC-01 cells may be prone to phenotypic changes in an *in vivo* environment (**Supplementary Fig. 1**).

### SarBC-01 organoids harbor genomic alterations which are shared with their parental tumor and representative of sarcomatoid malignancies

To further define the molecular profile of the SarBC-01 model, we performed whole exome sequencing (WES) analysis in the distinct components of the patient tumor (SARC and UroCa) and derived organoids at early and late passage (passage 6 and 20, respectively). Phylogenetic analysis based on WES highlighted a close relationship between all the analyzed samples, with 228 unique alterations shared by all specimens (Sar, UroCa, Organoids passage 6 and 20; **Fig. 3A** and **Supplementary Table 1**). Among these, we identified clonal somatic pathogenic mutations in genes that are frequently altered in sarcomatoid tumors from various epithelial entities including bladder, such as *TP53, RB1*, and *KRAS* (2,6) (**Fig. 3B, Supplementary Fig. 2**, and **Supplementary Table 1**). Additional common alterations included mutations and copy number changes in epigenetic regulators (e.g. *KMT2C, KMT2D)*, and in EMT-associated genes (e.g. *ZEB1, CDH1)* (**Fig. 3B**). In comparison, UroBC-01 organoids harbored pathogenic mutations in several bladder cancer-associated genes such as *TP53, TSC1*, and *ELF3*, which were also found in their parental tumor (**Supplementary Fig. 3, Supplementary Table 1**). Noteworthy, for both UroBC-01 and SarBC-01, main genomic drivers were clonal in all matched samples (i.e. tumor and organoids derived from same patient), while genomic subclones were detected at lesser cancer cell fractions in specific samples, highlighting intragenomic heterogeneity in tumor and organoid samples (**Supplementary Fig. 4**).

**Figure 3.**
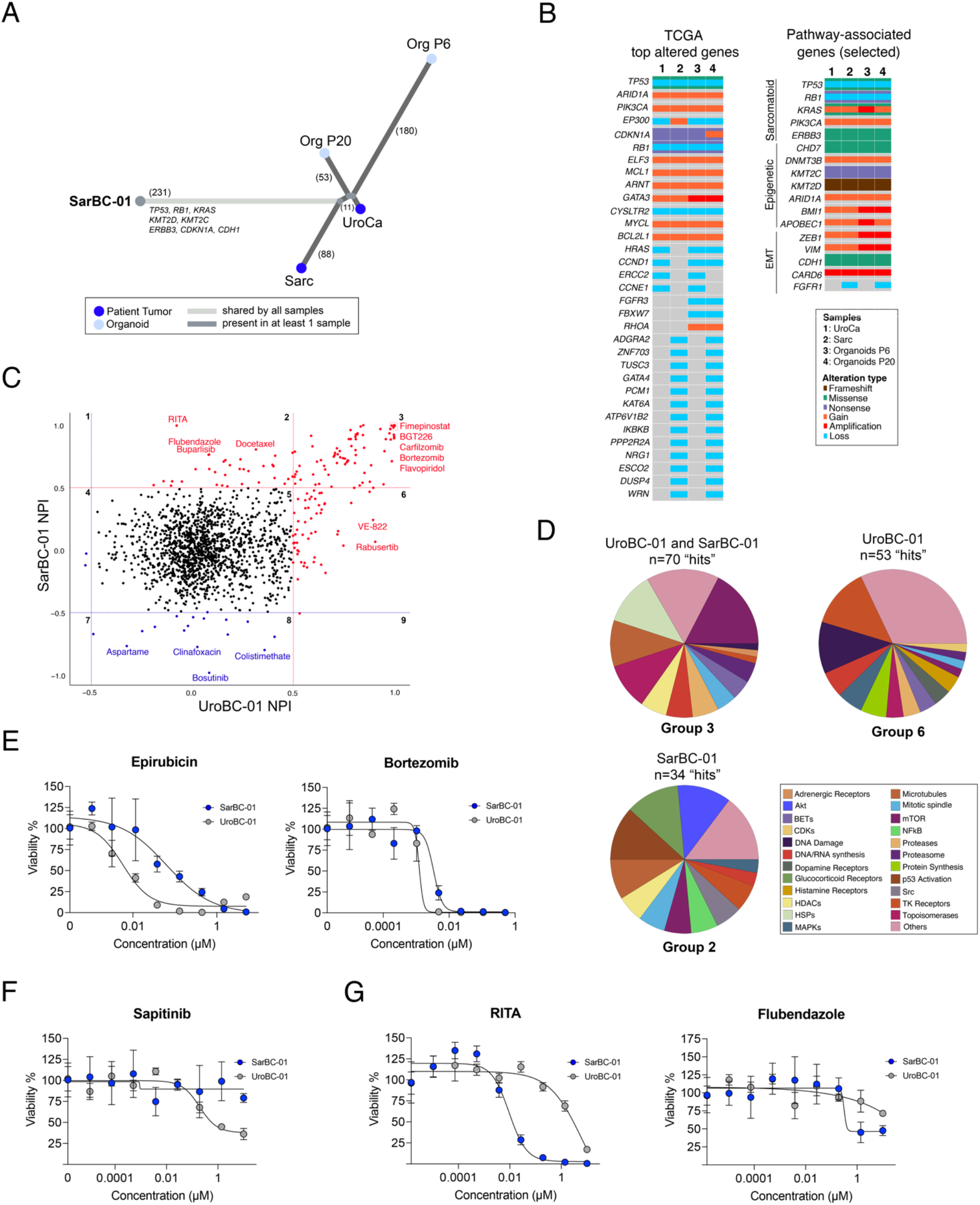
Identification of genomic drivers and drug sensitivities in SarBC-01 organoids. **(A)** Phylogenetic analysis of SarBC-01 patient tumors’ components and derived organoids at early and late passages using whole exome sequencing (WES). Numbers in brackets indicate the number of mutations. **(B)** Oncoplots depicting genomic alterations in the distinct SarBC-01 associated samples, as revealed by whole exome sequencing. Shown are alterations among the top 100 genes commonly mutated in BC (TCGA, **left panel**) and among selected genes associated with sarcomatoid cancers, epigenetic, and EMT pathways (**right panel**). Only alterations found in at least two samples are represented. A complete list of genomic alterations can be found in **Supplementary Table 1. (C)** Drug screening with a library of 1110 FDA-approved drugs and 457 clinical candidates identifies compounds with inhibitory effects (“hits”) in the Urothelial BC line (UroBC-01, group 6), the Sarcomatoid line (SarBC-01, group 2), and both lines (group 3), as well as compounds with “promoting” effects in each line (group 8). The dot plot shows normalized percentage inhibition (NPI) measured for each line, each dot representing one drug. “Hits” are defined as compounds associated with >50% inhibition as compared to negative controls (red dots, above the red line threshold). “Anti-hits” are defined as compounds associated with >50% growth promotion as compared to negative controls (blue dots, below the blue line threshold). **(D)** Pie charts summarizing the mode of action categories of drugs identified as hits for both UroBC-01 and SarBC-01 (top left, group 3), UroBC-01 only (top right, group 6), and SarBC-01only (bottom left, group 2). **(E-G)** Dose-responses curves for SarBC-01 and UroBC-01 organoids treated with a selected panel of drugs. Examples include drugs which are active in both lines **(E)**, only effective in UroBC-01 **(F)**, and more effective in SarBC-01 vs. UroBC-01 **(G)**. IC_50_ values for all tested compounds are reported in **Supplementary Table 3**.

Overall, these results highlight the close genomic relationship between UroCa and SARC concomitant components; importantly, identified genomic drivers are conserved in SarBC-01 organoids and representative of sarcomatoid malignancies.

### High-throughput screening identifies drug candidates targeting SarBC-01 and UroBC-01 organoids

Given that the SarBC-01 model displayed phenotypic and genomic features reminiscent of SARC, we sought to further exploit it to identify tailored therapeutic candidates. Following optimization (**Supplementary Fig. 5)**, we performed a high-throughput drug screening using a library comprising 1110 FDA-approved and 457 clinical compounds in both SarBC-01 and UroBC-01 (1 μM concentration in triplicates; see complete list of drugs in **Supplementary Table 2**). These analyses led to the identification of drug candidates inhibiting UroBC-01 cells exclusively (n=53 hits, group 6), or of SarBC-01 cells exclusively (n=34 hits, group 2), or of both (n=70 hits, group 3) (**Fig. 3C-D, Supplementary Table 2**). A subset of these drug candidates (n= 20), selected among standard-of-care and FDA-approved top hits in each group, was further tested at eight different concentrations to generate dose-response curves and determine the sensitivity profiles of UroBC-01 and SarBC-01 cells for each drug (**Fig. 3E-G, Supplementary Fig. 6**). Notably, although at different levels, both cell models were sensitive to chemotherapeutic drugs routinely used to treat bladder cancer (e.g. Cisplatin, Epirubicin, Gemcitabine); as an example, the half-maximal inhibitory concentration value (IC_50_) for Epirubicin was 10-fold higher in SarBC-01 cells as compared to UroBC-01 cells (IC_50=_ 0.063 μM vs. 0.006; see complete IC_50_ list in **Supplementary Table 3**). Clofarabine, a compound which was recently proposed to be highly effective in bladder cancer subtypes, was among the drugs inhibiting both UroBC-01 and SarBC-01 cells in our screen, albeit with medium efficacy in the SARC model (7) (**Supplementary Table 2**). Yet top hits for both models were enriched in compounds affecting the mTOR pathway, heat shock proteins, and microtubules (**Fig. 3C-E, Supplementary Fig. 6, Supplementary Table 2**). Interestingly, the most effective hits comprised several proteasome inhibitors such as Bortezomib (**Fig. 3E**), previously suggested to represent an attractive target in bladder cancer and other sarcomatoid tumors (8). Specific targeted compounds were found to be active in only one of the models, likely reflecting their unique molecular profiles. In particular, compounds targeting tyrosine kinase receptors and the MAP kinases cascade exclusively targeted urothelial UroBC-01 cells, while drugs affecting AKT and P53 pathways were specifically efficient against sarcomatoid SarBC-01 cells (**Fig. 3D-G, Supplementary Fig. 6**). Noteworthily, drugs with a mode of action linked to the glucocorticoid receptor (GR) pathway were not effective in the UroBC-01 model but were among the top categories effective in the SarBC-01 model (**Fig. 3D**). These drugs exclusively included GR agonists such as Prednisolone (Prdl) and Dexamethasone (Dex).

### High glucocorticoid receptor expression is a hallmark of SARC tumors

Prompted by this finding, we investigated the expression of GR in UroBC-01 and SarBC-01 tumors and derived models. Consistent with a specific response of SarBC-01 cells to GR-associated drugs, GR was highly expressed in SarBC-01 tumor cells; in contrast, its expression was restricted to stromal cells in the UroBC-01 tissue sample and absent in its derived tumor organoids (**Fig. 4A**). To assess whether high GR expression in SarBC-01 could be generalized to SARC bladder tumors, we analyzed a published dataset comprising transcriptomic data generated from 84 UroCa and 28 SARC samples (2). As expected, among the top genes positively enriched in UroCa were the gene encoding E-cadherin (*CDH1*) and other epithelial-associated genes such as *ELF3* and *TP63* (**Fig. 4B**). The top genes negatively enriched in UroCa included genes associated with EMT (e.g. *FGFR1, SNAI1, TGFB1*) as well as the gene encoding GR (*NR3C1*), which was significantly more expressed in SARC vs. UroCa samples (p < 0.0001, unpaired t-test; **Fig. 4B-C**). To validate these findings at the protein level and in another independent cohort, we assessed GR expression via immunohistochemistry in an in-house cohort comprising 14 UroCa and 13 SARC paraffin-embedded samples (**Supplementary Table 5**). In the UroCa group, GR expression was heterogenous with a large subset of low/negative tumors (n=8/14 samples with H-score< 50) and a minor subset exhibiting high expression (n=3/14 with H-score of 300). To note, two out of three UroCa samples with high GR expression were classified as “Basal/Squamous”. SARC tumor samples displayed significantly higher frequency of positivity (n=13/13 positive SARC) and higher H-score of GR, as compared to UroCa samples (p=0.02 Fisher’s exact test, p=0.03 Mann-Whitney test, respectively; **Fig. 4D, Supplementary Fig. 7, Supplementary Table 5**). In line with our human data, *nr3c1* was highly expressed in basal/squamous specimens and further higher in sarcomatoid samples, as compared to benign-like, hyperplasia, dysplasia and low-grade UroCa samples derived from a mouse model of bladder cancer (9).

**Figure 4.**
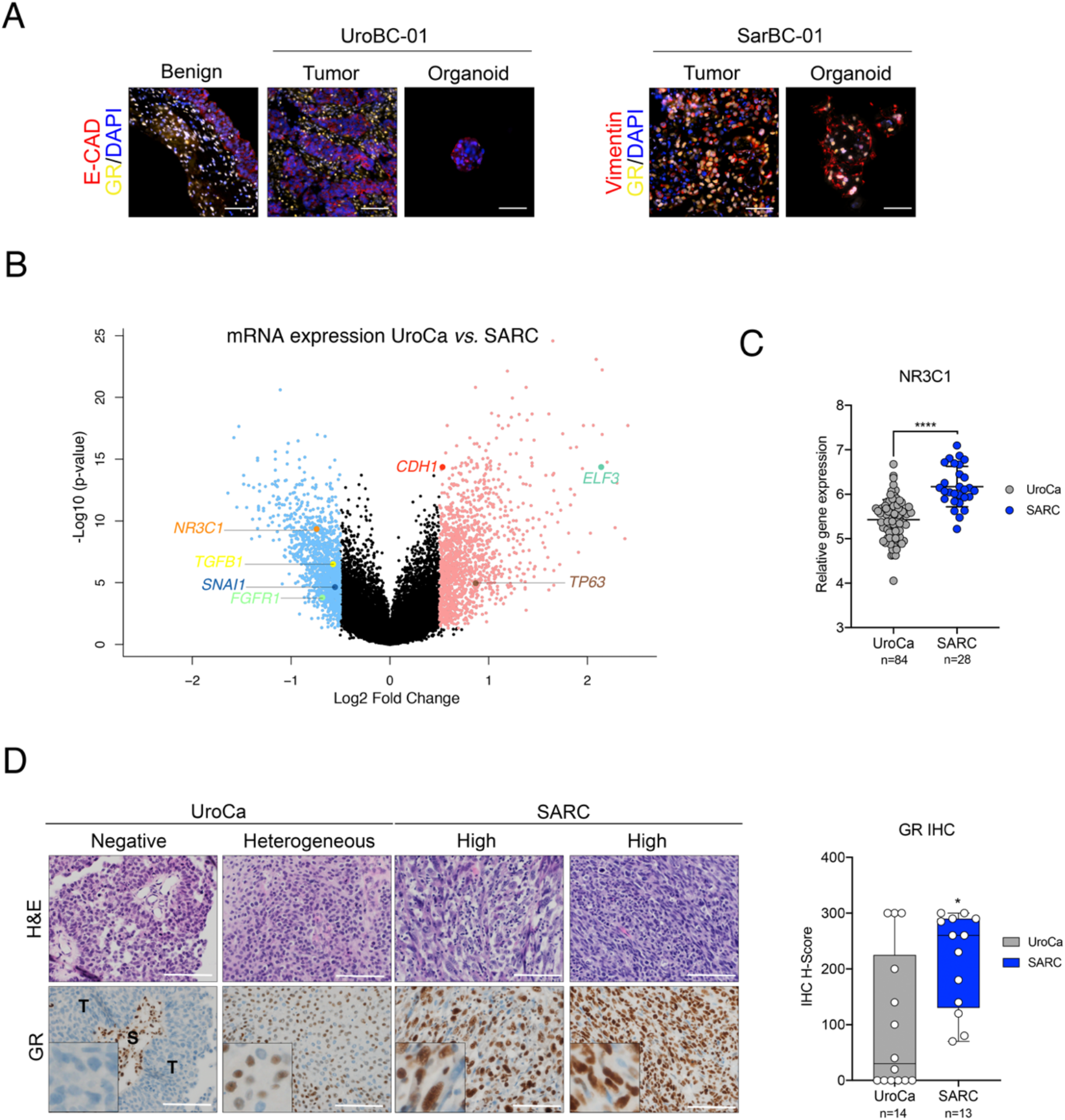
Glucocorticoid Receptor (GR) is highly expressed in SARC tumors. **(A)** Immunofluorescence analyses of UroBC-01 (left) and SarBC-01 (right) tumor and organoids pairs. Shown are representative images for the indicated antibodies. *GR: glucocorticoid receptor, E-cad: E-cadherin, Vim: Vimentin*. DAPI: 4’,6-diamidino-2-phenylindole. Scale bars represent 50 μm Volcano plot depicting differential gene expression between UroCa tumors (n=84) and SARC tumors (n=28) as analyzed from the dataset of Guo *et al*. (2). Indicated are selected genes significantly negatively or positively enriched in UroCa vs. SARC. **(C)** Expression of the *NR3C1* gene in UroCa and SARC samples. Statistical analysis was performed using an unpaired t-test (****: p < 0.001. **(D) Left panel:** Examples of H&E and immunohistochemical staining for GR in UroCa samples and SARC samples. *S: Stroma, T: tumor*. Scale bars represent 100 μm. **Right panel:** Box plot showing significant higher frequency of positivity and higher H-score for GR in SARC (n=13) vs. UroCa (n=14) samples (*: p < 0.05 for frequency and expression, Fisher’s exact test and Mann-Whitney test, respectively).

Thus, high GR expression is a feature characterizing a large majority of SARC tumors, which is well conserved in the SarBC-01 model.

### Glucocorticoid treatment leads to morphological and transcriptomic changes associated with EMT reversion

Prompted by these findings and the drug screen results, we next tested the effect of glucocorticoids (GC) in dose response experiments. Although their efficacy was mild in terms of inhibition of growth (∼70% of viability at a 1-10 μM concentration range), treatment with Prdl or Dex led to a notable change of morphology in SarBC-01 cells (**Fig. 5A-B**).

**Figure 5.**
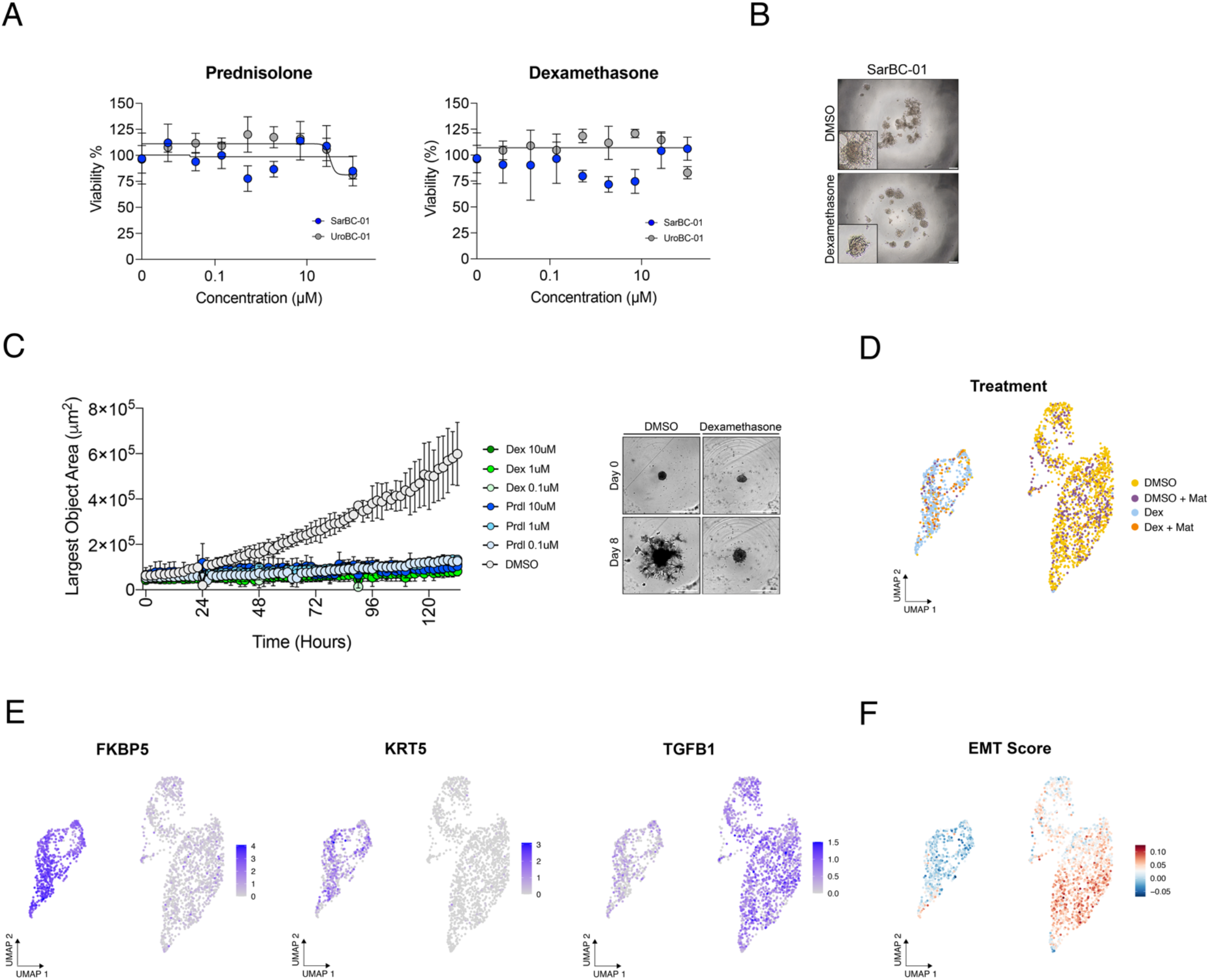
Glucocorticoid treatment leads to morphological and transcriptomic changes associated with reversion of epithelial-to-mesenchymal transition. **(A)** Glucocorticoids such as Prednisolone and Dexamethasone have a limited effect on the viability of SarBC-01 cells **(B)** Change of morphology of SarBC-01 upon treatment with 1 μM of Dexamethasone **(C) Left:** Largest bright field object area measured over time for spheroids generated from SarBC-01 cells treated with three concentrations of Dexamethasone (Dex) or Prednisolone (Prdl) as compared to the DMSO control. Two independent biological replicates were performed and one representative replicate is shown. **Right:** Representative images of SarBC-01 spheroids treated with DMSO or 0.1 μM of Dexamethasone at day 0 and day 8 are shown. Scale bars represent 800 μm. **(D-E)** Single-cell RNA sequencing of SarBC-01 cells following Dexamethasone treatment. Shown are UMAP representations of the treatment conditions **(D)** and expression of selected indicated genes **(E)**. 2170 cells were analyzed following incubation with DMSO or Dexamethasone in invasion assay conditions and in presence of absence of Matrigel, leading to 4 different culture conditions (DMSO, DMSO + Mat, Dex, Dex + Mat).

In line with these observations, the invasive ability of SarBC-01 cells was significantly reduced upon treatment with those drugs, even at low concentrations mirroring plasma levels in patients (0.1 μM Dex or Prdl vs. DMSO, mixed-effects analysis, p < 0.0001; **Fig. 5C**). To test whether GC-induced phenotypic effects on organoids correlated with transcriptomic changes at the cellular level, we performed single-cell RNA sequencing (scRNA-seq) in SarBC-01 cells treated with 0.1 μM Dex or with DMSO in invasion assay conditions. In order to account for potential effects of the Matrigel on the transcriptomic profile of the cells, the experiment was performed with and without Matrigel, leading to 4 distinct conditions (DMSO, DMSO + Matrigel, Dex, and Dex + Matrigel); prior to sequencing, these 4 conditions were multiplexed using MULTI-seq lipid-tagged indices, allowing to minimize technical confounders such as doublets and batch effects (10,11) (see **Material and Methods**). A total of 2170 cells was retrieved from the distinct conditions (median of 6700 genes per cell), which clustered based on treatment conditions and largely independently from the presence of Matrigel (**Fig. 5D**). Cells treated with Dex displayed significant higher levels of *FKBP5*, a GR target gene whose transcription is induced upon GR transactivation (12). Dex-treated cells were also characterized by gained expression of the two urothelial markers *KRT5* and *KRT7*, suggesting that SARC cells acquired epithelial-like transcriptomic features upon Dex treatment (**Fig. 5E, Supplementary Fig. 9**). Conversely, Dex-treated cells displayed significantly lower levels of genes associated with EMT such as *TGFB1, FGF2, SNAI2, IL1B, WNT5A, NOTCH1* and *VEGFA*, as compared to the DMSO condition (**Fig. 5E, Supplementary Fig. 9**). Consistent with these data, cells treated with Dex associated with a significantly lower EMT score as compared to cells treated with the DMSO control (**Fig. 5F**, score calculated based on a list of genes previously reported as EMT markers (13)).

Collectively, these data show that SARC cells gain epithelial-like features and lose mesenchymal-like features upon treatment with GC, suggesting that the SARC phenotype is plastic and may be therapeutically modulated.

## Discussion

A lack of experimental models emulating rare tumor entities, such as SARC, hampers the efforts to decipher mechanisms driving such diseases and the development of tailored clinical strategies. In this study, we address this limitation by establishing a 3D multicellular *in vitro* model derived from a SARC patient that retains key phenotypical and molecular features of SARC and is tumorigenic *in vivo*. While few SARC-like models have recently been described (7,14), SarBC-01 represents the first fully characterized long-term organoid model derived from a SARC patient; such type of model better recapitulates heterogeneity than cell lines and is more easily amenable to drug screen than xenografts, making it an attractive model for basic and translational research.

High-throughput drug screening highlighted the efficacy of standard chemotherapies, as exemplified by Cisplatin which was highly active in SARC cells. Further, these analyses revealed potential novel therapeutic options displaying higher efficacy as compared to the standard of care, such as the proteasome inhibitor Bortezomib and the p53 interacting molecule RITA. Importantly, the SarBC-01 model is derived from one single patient and we acknowledge that the identification of specific targeted compounds may reflect its unique genomic profile rather than its sarcomatoid identity. Nevertheless, SarBC-01 cells harbor mutations in genes that are commonly altered in a broad range of SARC tumors including but not limited to the bladder context (2,6) as well as phenotypical features typical of SARC, making it a model that could be broadly generalized to sarcomatoid tumors.

Notably, drugs with a mode of action linked to the glucocorticoid receptor (GR) pathway were among the top categories effective in the SarBC-01 model. These data led us to investigate the expression of the steroid hormone receptor GR in tumors samples obtained from UroCa and SARC. Exploiting two distinct cohorts, we showed that high *NR3C1*/GR expression, at the mRNA and protein level, is a feature characterizing a large subset of SARC tumors. While GR signaling has not been extensively studied in the context of bladder cancer, limited available data have pointed to a multifaceted and context-specific role of this pathway (12,15,16). In UroCa, GR expression has been shown to be lower in high-grade *vs*. low-grade as well as in muscle-invasive *vs*. non-muscle-invasive disease, consistent with a potential tumor-suppressor function (12). Noteworthily, while we observed a large subset of muscle-invasive UroCa tumors with negative GR, two out of three UroCa samples displaying high GR expression were classified as “Basal/Squamous”, a subtype from which SARC has been proposed to originate from (2). In contrast to UroCa, the relevance of GR and its associated pathway had not been addressed in the context of histological variants of bladder cancer, such as SARC so far. Noteworthily, other sarcomatoid tumor entities have been suggested to display high GR levels, suggesting that our findings may have relevance beyond the bladder context (17).

Due to high GR levels, SARC tumors may be particularly sensitive to GR signaling-associated compounds such as GC, as compared to most UroCa tumors. In agreement with this hypothesis, we show that GC treatment induces a limited decrease of viability, as well as epithelial-like morphological and transcriptomic changes which are consistent with (partial) reversion of EMT in SARC cells (*See hypothetical model in* **Fig. S10**). These data therefore suggest a potential benefit of GC compounds which are commonly used in an adjuvant and palliative setting in various tumor types (18) and warrant future research to investigate mechanisms driving their effects. Given the complex role of GR signaling in bladder carcinogenesis, a precise assessment of the histological subtype and of GR expression in patients’ tumor samples may allow informed decision regarding their use.

Collectively, our data highlight the high plasticity potential of the SARC phenotype. We first show that the SARC tumor and its derived cells display a high degree of genomic similarity with their UroCa counterpart, suggesting a common ancestor as previously proposed (2,3). The absence of relevant private genomic alterations in the SARC counterpart potentially explaining the evolution of the cells from UroCa to SARC and the partial recovery of epithelial features in xenografts, raise the hypothesis that SARC differentiation might be a reversible phenomenon driven by microenvironmental cues. Finally, the EMT reversal effects associated with GC treatment further emphasize the plasticity of the SARC phenotype and suggest that it may be therapeutically modulated towards less aggressive phenotypes.

Altogether, our study highlights the power of organoid models to identify precision oncology strategies for rare tumor entities and to provide key insights into factors driving these diseases.

## Material and Methods

### Patients and clinical samples

Bladder cancer samples used for organoid generation were obtained from two patients operated at the University Hospital of Basel (USB) under an approval by the Ethical Committee of Northwestern and Central Switzerland (EKBB 37/13). Clinical and pathological characteristics of the patients and associated samples are described below:

| Patient | Sex | Age | Smoker | Previous intervention | Stage and Grade | Sample | Histology |
| --- | --- | --- | --- | --- | --- | --- | --- |
| SarBC-01 | Male | 73 | yes | none | ≥ pT2, G3 | TUR-B | UroCa with SARC components |
| UroBC-01 | Male | 75 | yes | none | pT2, G3 | Cystectomy | Poorly differentiated UroCa |

In addition, immunohistochemical analysis for GR expression was performed on an archival cohort of 14 UroCa samples and 13 SARC samples under an approval by the Ethical Committee of Northwestern and Central Switzerland (EKNZ 2014-313). Clinical and pathological characteristics of this cohort is described in **Table S5**.

### *In vitro* model generation

Bladder cancer tissues were washed in aDMEM/F12 (GIBCO). One portion of the sample was fixed in formalin 4% and kept at 4°C until further processing. The rest of the sample was processed as described in (4). Briefly, the tissue was mechanically and enzymatically digested, and filtered through a 100 μm strainer (Corning) to generate a cell suspension. Cells were counted and resuspended in order to reach a concentration of 5×10^3^ cells/well in a 70/30 ratio Matrigel (Corning)/organoid culture medium. The organoid culture medium was comprised of aDMEM/F12 supplemented with 100 ng/ml WNT3A (Bio-Techne), 100 ng/ml R-Spondin1 (R&D Systems), 50 ng/ml EGF (Peprotech), 1mM N-acetyl-L-cysteine (Thermo Fisher), 100 ng/ml Noggin (Peprotech), 1 μM TGFβ inhibitor (Selleck), 1x N2 (GIBCO), 1x B27 (GIBCO)) solution (adapted from (19)). Subsequently, cells were plated in a 10 μL drop at the bottom of the well of a 96-wells plate and incubated at 37°C and 5% CO2 for 30 minutes (min), resulting in the embedment of the cells in a solid Matrigel drop (embedded condition; EMB). 100 μL of organoid culture medium was added to the wells and fresh medium was dispensed every 3-4 days; organoids were maintained for a maximum of 30 days in culture before passaging at a 1:2 dilution. For passaging, Matrigel was digested by adding 1 mg/ml dispase (GIBCO) solution to the wells and incubating for 60 min at 37°C before harvesting. The collected organoids underwent enzymatical digestion to generate a suspension of cells, that were re-seeded in EMB conditions (for expansion) or resuspended in culture medium containing 5% Matrigel and plated in low-adherence 96 well plates for all the subsequent analysis (non-embedded condition; NE). All *in vitro* and *in vivo* analyses were performed in NE conditions.

### Histology, immunohistochemistry and immunofluorescence

Histological analysis was performed by standard hematoxylin and eosin (H&E) staining. Immunohistochemical analyses were conducted according to standard indirect immunoperoxidase procedures. Immunofluorescence staining was performed on 4-μm-thick sections, following antigen retrieval with boiling citrate acid-based antigen masking solution at 98°C for 15 min such as in (20). Immunofluorescence images were captured using a Nikon Ti2 microscope. Details of all the antibodies and dilutions are provided in **Supplementary Table 4**.

### Invasion assay

The invasion assay was performed as described in (21). Briefly, single cells derived from organoids were plated in low-adherence U-shaped 96-wells plates at a concentration of 1000, 500, 250, 125 cells/well in organoid culture medium. After 8 days in culture, spheroids formed at the bottom of the wells, and half of the culture medium was carefully removed and replaced with ice-cold Matrigel. The plates were spun down at 350g for 10 min at 4°C and placed in the Incucyte® Live-Cell Analysis System (Sartorius). The growth of the spheroids was monitored for the following 8 days and invasion was automatically quantified via the Incucyte® Spheroid module analysis tool, measuring the increase of the spheroids area over time.

To assess the invasion capacity upon glucocorticoids treatment, a similar approach was applied. Briefly, 1000 cells were plated in low-adherence U-shaped 96-wells plates and supplemented with culture medium containing Dexamethasone 10 μM, 1 μM, 0.1 μM, Prednisolone10 μM, 1 μM, 0.1 μM or 1% DMSO as negative control in triplicates. After 8 days in culture, half of the culture medium was carefully removed and replaced with ice-cold Matrigel. The spheroid growth was monitored via Incucyte® for the following 8 days and quantified with the Spheroid module analysis tool. The invasion capacity was assessed at passage 40 and 50 of SarBC-01 and passage 39 of UroBC-01.

### *In vivo* tumorigenic capacity

All mouse experiments were conducted with the approval of the Animal Care Committee of the Kanton Basel-Stadt, Switzerland (3066-32428). Mice were bred and maintained in the animal facility of the Department of Biomedicine of the University Hospital Basel under specific pathogen-free conditions on a 12h day and 12h night schedule with *ab libitum* access to food and drinking water. Single cell suspensions derived from organoids were spun down and resuspended at a 50/50 ratio Matrigel/PBS solution. One million of cells was subcutaneously injected in the flank of NOD *scid* gamma (NSG); two mice were injected per organoid line to compare the tumorigenicity capacity between the two lines (passage 32 and 31 for SarBC-01 and UroBC-01, respectively). In addition, 2 mice were injected to confirm the tumorigenic potential of SarBC-01 cells at a different passage (passage 36, 2/2 mice with tumors). Tumor size was measured twice weekly and tumors were harvested when reached 1500 mm^3^ in volume. The human origin of all patient-derived organoid xenografts (PDOXs) was confirmed by staining with an antibody recognizing human mitochondria (ab92824, Abcam).

### DNA extraction

For SarBC-01, DNA was extracted from formalin-fixed paraffin-embedded (FFPE) material (SARC and UroCa, components; healthy lymph nodes for germline DNA), and flash frozen organoids. For UroBC-01, DNA was extracted from fresh peripheral blood mononucleated cells (PBMCs) for germline and from flash frozen material for the parental tumor and derived organoids. For FFPE tissue, 10-μm thick unstained tissue sections were cut on glass slides. The distinct morphological components of sarcomatoid and urothelial bladder cancer (SARC and UroCa respectively) were identified and marked by a trained pathologist. The samples were deparaffinized with incubation in Xylene for 5 min and the regions of interest scratched from the glass slide and collected. DNA was extracted using the RecoverAll RNA/DNA extraction kit (Invitrogen, Carlsbad, CA, USA), according to the manufacturer’s instructions. The collected DNA underwent incubation with Uracil-DNA Glycosylase at 37°C for an hour, followed by an incubation at 50°C for 10 min. For fresh and flash frozen tissue, DNA and RNA were simultaneously isolated using the Quick-DNA/RNA Miniprep kit (Zymo Research, Irvine, CA, USA), according to the manufacturer’s instructions. Flash frozen tissues were crushed in liquid nitrogen with plastic pestels (Fisher, Thermo Fisher Scientific, Inc., Waltham, MA, USA) prior to isolation. DNA were quantified using the Qubit Fluorimeter assay (Thermo Fisher Scientific).

### Whole exome sequencing (WES) and variant annotation

DNA extracted from FFPE and flash frozen specimens was subjected to WES. Twist Human Core Exome + RefSeq + Mito-Panel kit (Twist Bioscience) was used for the whole exome capturing according to manufacturer’s guidelines. Sequencing was performed on Illumina NovaSeq 6000 using paired-end 100-bp reads and yielded a mean depth of coverage comprised between 108.6x and 153.4x. Sequencing was performed by CeGaT (Tübingen, Germany) and FASTQ processing workflow was adapted from (22). Briefly, reads were aligned to the reference human genome GRCh38. Somatic variants were detected using MuTect2 (23) and discarded if they had a variant allelic fraction < 5% or if were covered by fewer than 3 reads. To filter out potential artifacts, we further excluded variants present in more than two of a panel of 123 non-tumor samples. The complete list of identified variants is provided in **Supplementary Table 1**. Variant annotation was performed by SnpEff software v.4.1 (24). The heatmap of non-synonymous mutations was generated using the R package maftools v.2.10.5 (25).

### Copy number aberration and clonal analysis

Allele-specific CNAs were identified using FACETS v.0.5.6 (26). Genes were classified using the following criteria: “gains”: total copy number greater than gene-level median ploidy; “amplification”: total copy number greater than ploidy +4; losses: total copy number less than ploidy; homozygous deletion: total copy number of 0. Loss of heterozygosity were identified as those where the lesser (minor) copy number state at the locus was 0. Clonal analysis was performed with the ABSOLUTE V2.0 (27) algorithm. Solutions from ABSOLUTE were manually curated to assure the solution matched the ploidy estimate generated by FACETS. For heatmap representations, mutations and copy number alterations were shown based on top 100 altered genes in muscle-invasive bladder cancer (TCGA study (28)), or selected genes associated with specific pathways (sarcomatoid (6), epigenetic (29), EMT (13)).

### Phylogenetic analysis

A neighbor joining tree was built for each case based on the repertoire of non-synonymous and synonymous somatic mutations and small deletions, as described by (30). A mutation that was found in all samples deriving from one patient was considered as ‘trunk’. Mutations that were detected in only one of the samples from the same patient were classified as ‘private’. Furthermore, cancer cell fraction estimates generated by ABSOLUTE were used as input to PhylogicNDT (31) to find mutation clusters, infer subclonal populations and their phylogenetic relationships.

### Drug screening

To evaluate the assay quality, wells were plated in NE conditions at a concentration of 250, 500, or 1000 cells/well in low-adherence 384 well plates (Greiner Bio-One, St.Gallen, Switzerland). After 5 days, 12.5 μL of organoids culture medium containing either DMSO (negative control) or staurosporine (positive control) was added and after 5 additional days, the viability of the cells was assessed by luminescence using CellTiter-Glo 3D (Promega) according to manufacturer instructions. Luminescence was read on a Synergy H1 Multi-Mode Reader (BioTek Instruments). The z’ factor for each seeding concentration was calculated as:

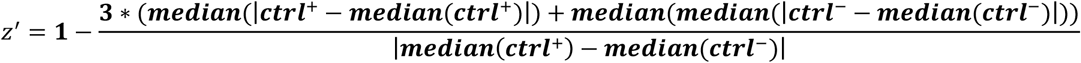

An assay was considered of good quality if it had a z’ factor above 0 and no positional effects. For the main assay, cells were plated at a concentration of 1000 cells/well. 5 days after plating, 1567 compounds (1110 FDA approved drugs and 457 in clinical trial drugs, obtained from NEXUS Personalized Health Technologies, Zürich, Switzerland) were added in triplicates at a 1 μM concentration. After 5 days of treatment, the viability of the organoids was measured as described above and normalized to the negative control); The normalized percentage inhibition (NPI) was calculated for each drug as:

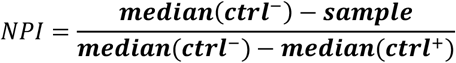

Compounds were defined as “hits”, if the median of the NPI was higher than 0.5. Compounds were defined as “anti-hits”, if the median of the NPI was lower than -0.5 (**Supplementary Table 2**).

### Drug-dose response analysis

To validate results from the drug screen, cells were plated at a concentration of 1000 cells/ well in low-adherence 384 well plates. 20 drugs from the identified hits were selected and an 8-point dilution series of each compound was dispensed in triplicate using a Tecan Digital Dispenser D300e (Tecan). Drug concentrations spanned from 10 pM to 1 mM, depending on the drug. Cell viability was measured by CellTiter-Glo 3D assay following 5 days of drug incubation, and results were normalized to untreated controls. Data analyses were performed using GraphPad Prism, and the values of IC_50_ (see **Supplementary Table 3)**, Hill slope, and AUC were calculated by applying nonlinear regression (curve fit) and the equation log(inhibitor) vs. normalized response (variable slope).

### In silico mRNA expression analysis

Public datasets were downloaded from the GEO database (Human dataset: accession number <u>GSE128192; mice dataset: accession number GSE197016</u>). The human dataset comprised 28 cases of SARC and 84 cases of UroCa (2). Differential gene expression analysis on this dataset was performed using the GEO2R online tool with default parameters. In brief, comparison between SARC and UroCa samples was carried out using the limma package and the p-value were adjusted using the Benjamini-Hochberg correction. Genes with FDR–adjusted p values < 0.05 and fold changes > 0.5 were considered as differentially expressed. The comparison of *NR3C1* expression levels was carried out using an unpaired t-test. The mouse dataset comprised 37 bladder cancer samples deriving from mice exposed to N-butyl-N-(4-hydroxybutyl)-nitrosamine (BBN) (9). The data were log-normalized and the samples classified and ordered based on their pathological stage, as described in the original publication. *Nr3c1* expression levels were plotted for each sample class.

### GR protein expression analysis

The antibody against GR (ab183127, Abcam) was tested on sections from formalin-fixed paraffin-embedded kidney, testis, and pancreas positive controls and assessed by a trained pathologist. A composite scoring system (H-score) was used by multiplying a given nuclear intensity (between 0 and 3) by the percentage of positive cells. Details of the patients’ cohorts and staining results can be found in **Table S5**.

### Single cell RNA-sequencing (scRNA-seq) sample preparation and multiplexing

SarBC-01 cells derived from organoids were plated in low-adherence U-shaped 96-wells plates at a concentration of 2000 cells/well and supplemented with medium containing either 100 nM Dexamethasone (Dex) or 1% DMSO as negative control. After 8 days in culture, spheroids formed at the bottom of the wells, and half of the culture medium was carefully removed and replaced either with ice-cold Matrigel (Mat) or freshly prepared medium, resulting in 4 different culture conditions (+Mat/Dex, +Mat/DMSO, -Mat/Dex, -Mat/DMSO). After 8 days, the spheroids were harvested upon incubation with TrypLe for 15 min, dissociated to single cells by gentle pipetting and washed in PBS. Cells belonging to the distinct culture conditions were subsequently processed for multiplexing using the MULTI-seq protocol (10). Briefly, a mix of lipid-modified DNA oligonucleotides and unique barcode oligonucleotides for each culture condition was added to the cells and incubated in cold PBS for 5 min. Next, a lipid-modified co-anchor was added to each sample to stabilize the membrane bound barcodes. After a 5-min incubation on ice, cells were washed in PBS containing 1% FBS 1% BSA to quench unbound barcodes. Cell number and cell viability was then assessed for each sample with an automatic hematocytometer. Finally, samples were pooled together with comparable cell number, washed with PBS 1% FBS 1% BSA, and 15 000 cells were loaded in a Chromium Single Cell 3′ GEM Library and Gel Bead Kit v3 (10x Genomics).

### scRNA-seq library preparation, sequencing, and quality control

Gene expression (cDNA) and MULTI-seq libraries were prepared according to the manufacturers’ protocol of Chromium Next GEM Single Cell 3’ reagents Kits v3.1 (Dual Index). In brief, after GEMs embedding, cDNA was generated by reverse transcription reactions. MULTI-seq barcode fragments were separated from endogenous cDNA fragments during the first round of size selection using SPRIselect beads (Beckman coulter). Next, cDNA and MULTI-seq fragments were processed separately. Fragmentation, end repair, and A-tailing procedure were performed on endogenous cDNA fragments, and sample dual indexes (Dual Index TT Set A plate) were lastly added over PCa amplification. After clean-up, MULTI-seq barcode fragments were PCR amplified and tagged with RPI index (TruSeq technology) and i5 universal index. Both libraries were cleaned up with SPRIselect beads to avoid primer contamination and fragments with inappropriate size (10). cDNA and MULTI-seq libraries were analyzed using an Agilent Bioanalyzer (DNA High Sensitivity kit) prior to sequencing on a NovaSeq6000 (S2 flow cell) platform. Cellranger (v.6.1.2) was used to process the cDNA raw data and generate a raw count matrix that was subsequently loaded in the R package Seurat (v.4.3.0) (32). Sample barcodes were demultiplexed using the HTODemux function implemented in Seurat (10). Cells labelled as doublet or negative were removed from the Seurat object. Furthermore, cells with less than 2500 detected genes and more than 25% mitochondrial transcripts were filtered out. A total of 2170 cells was obtained and subsequently analyzed.

### scRNA-seq bioinformatic analysis

The Seurat dataset was log normalized and 2000 variable features were initially computed using the ‘vst’ method. Data were scaled and the first 30 principal components were calculated for downstream analyses. Uniform Manifold Approximation and Projection (UMAP)(33) used for visualization. Cell clusters that were including the cells within the different culture conditions were identified with the ‘FindClusters’ function offered by Seurat with a resolution of 0.8. Exploratory analysis was performed to visualize the expression of genes linked to glucocorticoid receptor pathway and EMT. The EMT score was calculated with the ‘AddModuleScore’ function, using a list of genes previously reported as EMT markers (13). Differential gene expression analysis was performed with the MAST method (34) implemented in Seurat within the ‘FindMarkers’ function. The analysis was limited to genes detected in at least 25% of the tested populations and showing at least ±0.5 log fold change difference.

### Statistical analysis and data representation

The Prism GraphPad software (version 8.0, San Diego, CA, USA) was used for statistical analyses and data representation. Statistical analysis of invasion assays was conducted by mixed-levels analysis. The comparison of *NR3C1* mRNA levels between UroCa and SARC was conducted by an unpaired t-test. For GR protein expression, the average of the H-Score was compared with a Mann-Whitney test, while the frequency of positivity in each group was assessed using a Fisher’s exact test. For all statistical analyses, p values <0.05 were considered significant.

### Data availability

The whole exome sequencing and the single-cell RNA sequencing data generated in this paper have been submitted to the European Genome-phenome Archive. Before publication, data will be available upon request to the corresponding author of the study.

## Acknowledgments

We thank the patients who consented to participate to the study and all clinicians, pathologists, and study nurses involved in the enrollment of patients in the study and in the samples acquisition. We are grateful to Ilaria Aborelli for experimental help and to Raphaelle Servant, Mattia Marinucci, and Jan Schneeberger for support with the animal experiments. We acknowledge the assistance of the IHC facility of the Institute of Pathology, the microscopy and bioinformatics core facilities of the DBM, and Tijmen Booij of NEXUS Personalized Health Technologies (ETH Zurich, Switzerland) for providing the compound libraries; we thank the Next Generation Sequencing Platform of the University of Bern for performing the scRNA sequencing. We thank Renaud Mével for laying the groundwork for multiplexed scRNA-seq analysis and Christopher McGinnis and Zev Gartner (UCSF, US) for providing MULTI-seq reagents.

Financial support was provided by funding from the Swiss Cancer Research Foundation (KFS-4983-02-2020 to CL) and the Department of Surgery of the University Hospital Basel (*PMC* Platform).

## Authors contributions

Conceptualization: M.G., L.B., S.N., R.P., C.L.M.; Validation: M.G.; Formal analysis: M.G., L.B., S.N., L.R., R.P.; Resources: L.B., S.N., M.C., H.P., T.V., H.S., C.A.R., L.B.; Investigation: M.G.; Software: L.R.; Data Curation: L.R., F.S., C.A.R, L.B.; Visualization: C.L.M.; Writing - Original Draft: M.G., C.L.M.; Writing - Review & Editing: M.G., L.B., S.N., R.P., L.R., M.C., H.P., S.P., T.V., F.S., H.S., C.A.R., L.B., C.L.M.; Supervision: C.L.M.; Funding acquisition: C.L.M.

