## Supplementary Material for "Patient-derived organoids identify tailored therapeutic options and determinants of plasticity in sarcomatoid urothelial bladder cancer"

### Supplementary Material List

#### Supplementary Figures

Figure S1. Characterization of SarBC-01 and UroBC-01 Patient-Derived Organoid Xenografts (PDOXs)

Figure S2. Frequency of specific genomic alterations in sarcomatoid tumors

Figure S3. Whole Exome Sequencing Analysis of UroBC-01 tumor and matched organoids

Figure S4. Clonal evolution in SarBC-01 and UroBC-01 tumors and derived models

Figure S5. Optimization and quality control of the high-throughput drug screen

Figure S6. Drug response analysis for selected compounds

Figure S7. Glucocorticoid Receptor (GR) expression in UroCa and SARC samples

Figure S8. Expression of *Nr3c1* in a mouse model of bladder cancer progression

Figure S9. Single-cell RNA sequencing of SarBC-01 cells following Dexamethasone treatment

Figure S10. Hypothetical model: Glucocorticoid Receptor signaling in SARC tumors

#### Supplementary Tables

Table S1. List of somatic mutations (excel file)

Table S2. Hight throughput drug screen (excel file)

Table S3. IC50 of selected compounds (excel file)

Table S4. List of antibodies used in this study (excel file)

Tabel S5. Patients’ cohort for GR immunohistochemical analysis

#### Supplementary Figures

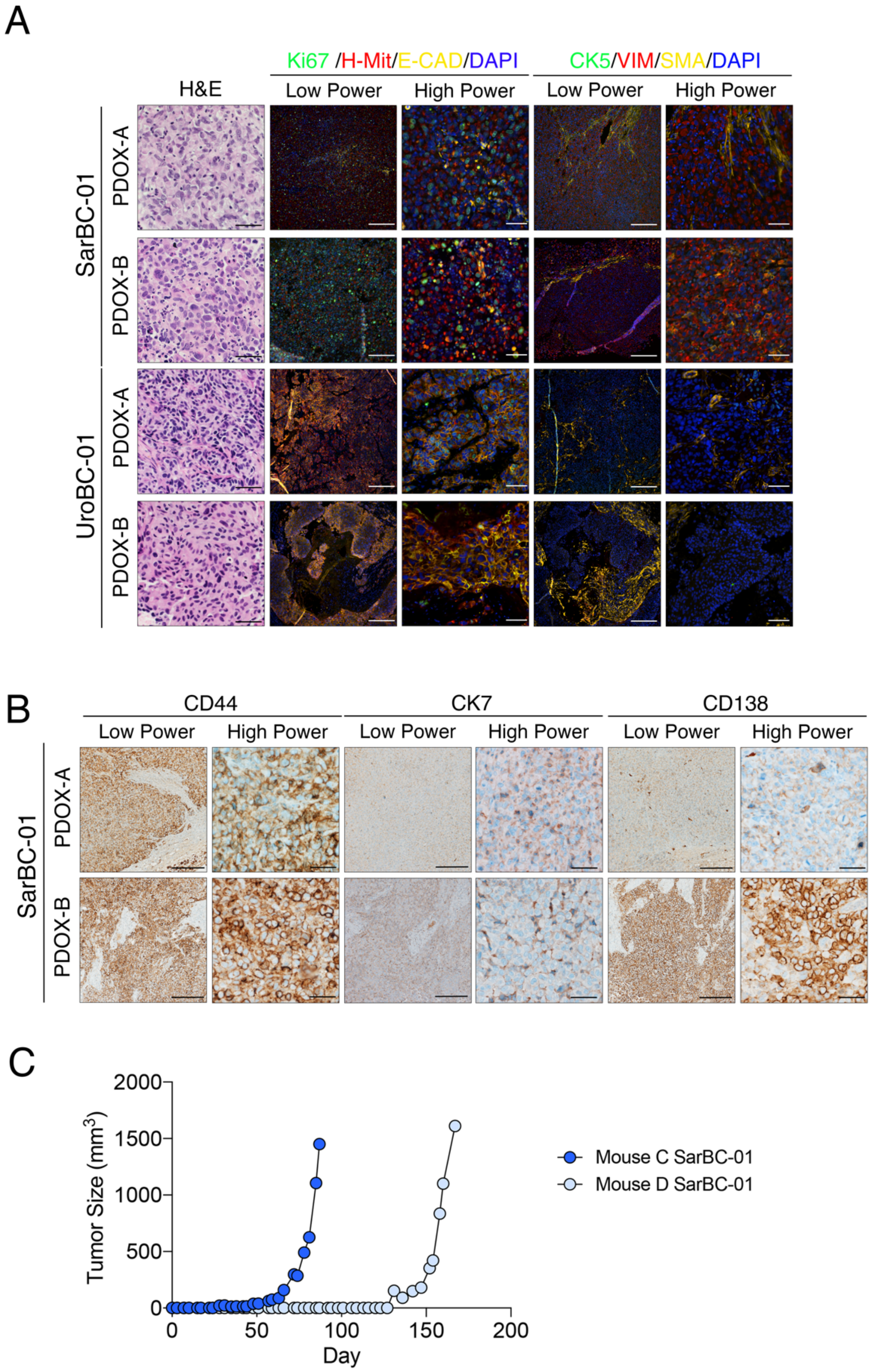


**Figure S1. Characterization of SarBC-01 and UroBC-01 Patient-Derived Organoid Xenografts (PDOXs). (A)** H&E and immunofluorescence analyses of SarBC-01 (top) and UroBC-01 (bottom) PDOXs. Shown are representative images for the indicated antibodies. *H-*

*Mit: Human mitochondria, Vim: Vimentin, E-cad: E-cadherin, SMA: Smooth Muscle Actin. DAPI: 4',6-diamidino-2-phenylindole.* Scale bars represent 50 μm for high magnification images and 250 μm for low magnification images. **(B)** Representative IHC staining for the indicated antibodies, showing expression of specific epithelial markers in the xenografts derived from SarBC-01 cells. Scale bars represent 50 μm for high magnification images and 250 μm for low magnification images. **(C)** Growth kinetic of xenografts generated by sucutaneus inhection of SarBC-01 cells in two additional NGS mice (PDOX C and D).

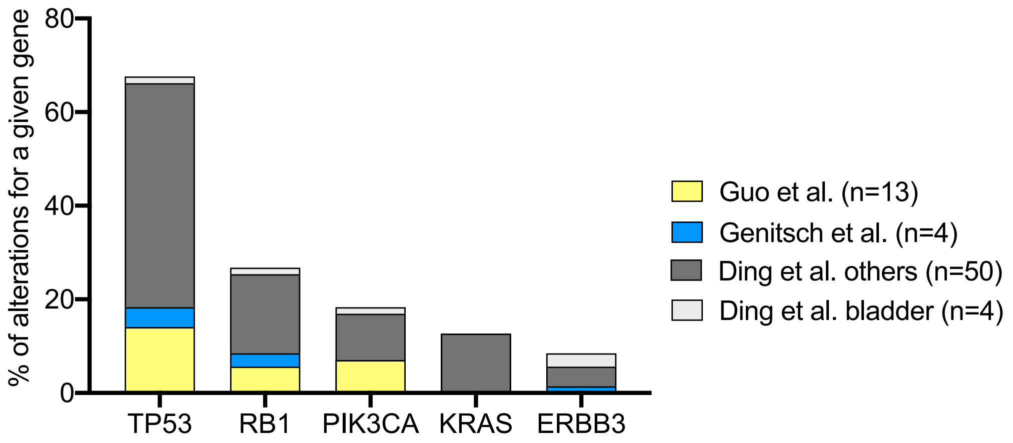


**Figure S2. Frequency of specific genomic alterations in sarcomatoid tumors.** Four datasets were included in the analysis, integrating genomic data obtained from sarcomatoid bladder tumors (*Guo et al., Genitsch et al., Ding et al. bladder*) as well as other sarcomatoid tumor entities (*Ding et al., others*).

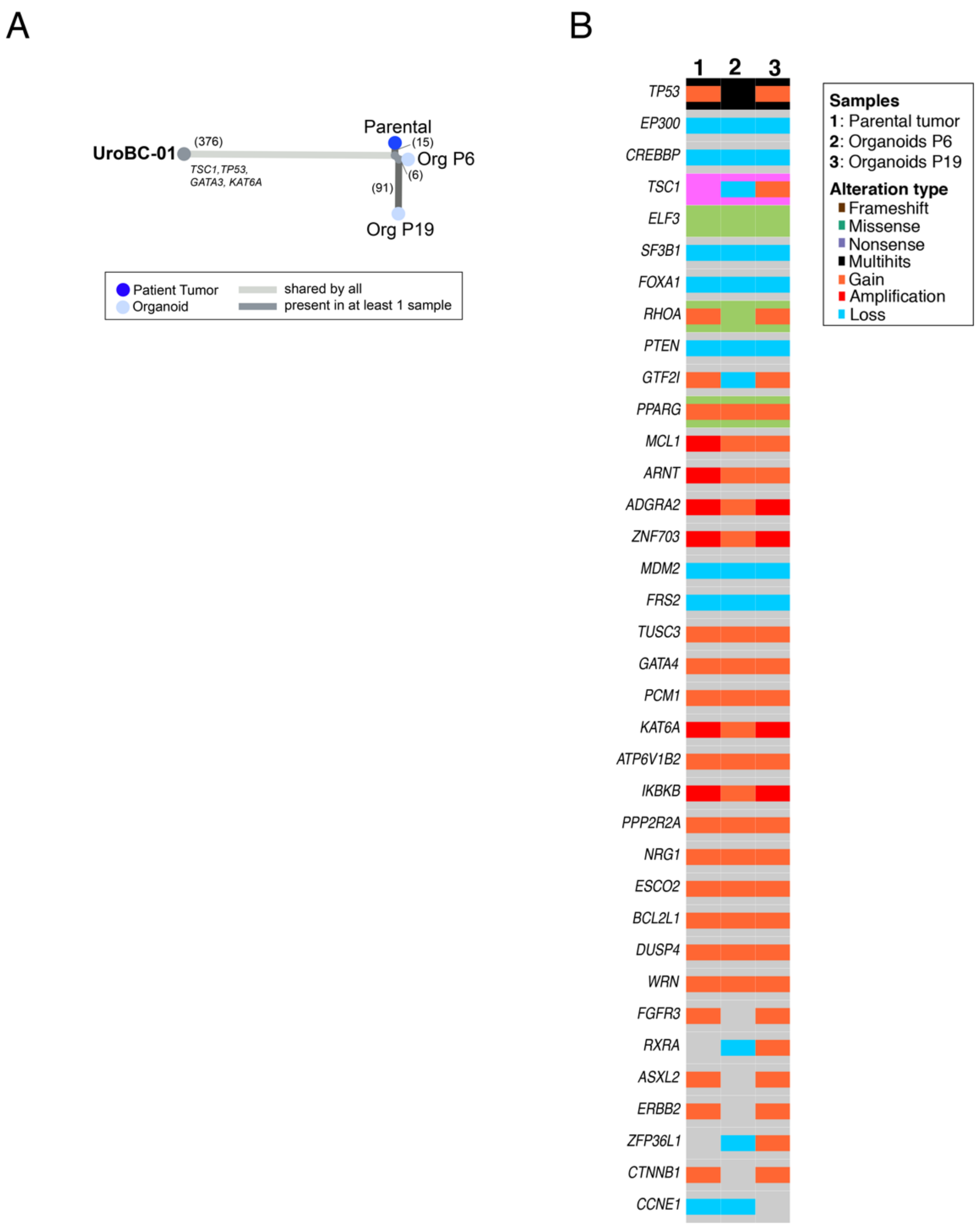


**Figure S3. Whole Exome Sequencing Analysis of UroBC-01 tumor and matched organoids. (A)** Phylogenetic analysis of UroBC-01 patient tumor and derived organoids at early and late passages (passage 6 and 19, respectively) using whole exome sequencing (WES). Numbers in brackets indicate the number of mutations **(B)** Oncoplots depicting genomic alterations in UroBC-01 tumor and its derived organoids, as revealed by whole exome sequencing. Shown are alterations among the top 100 genes commonly mutated in BC (TCGA). Only alterations found in at least two samples are represented. A complete list of genomic alterations can be found in **Supplementary Table 1**.

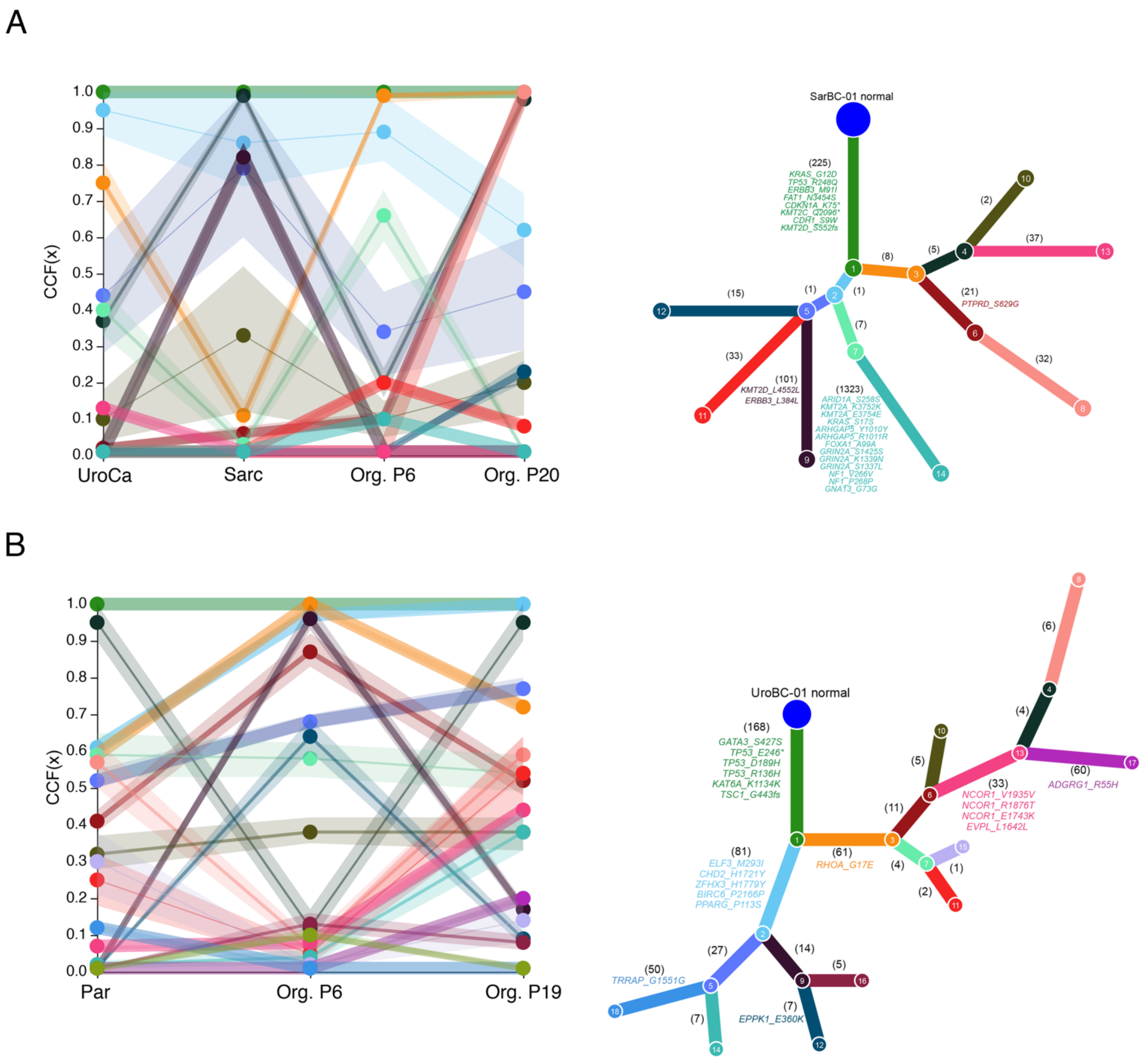


**Figure S4. Clonal evolution in SarBC-01 and UroBC-01 tumors and derived models. (A)** UroCa tumor, Sarc tumor, and its derived organoids at passage 6 (Org. P6) and passage 20 (Org. P20) for patient SarBC-01. **(B)** Parental tumor (Par) and its derived organoids at passage 6 (Org. P6) and passage 19 (Org. P19) for patient UroBC-01. For both **(A)** and **(B)**, trace plots (left panels) show cancer cell fraction (CCF) for each mutational cluster in each sample. Ribbons show 95% confidence intervals, while center of bands show mean CCF estimate. Phylogenetic trees (right panels) show best solution for evolutionary relationship between clones with different clusters of mutations where each node (numbered) is a cluster of mutations. Numbers on each branch show the number of mutations distinguishing a clone from the previous clone (all genes). Potential driver genes mutations distinguishing between a clone and the previous are indicated in colors corresponding to the branch.

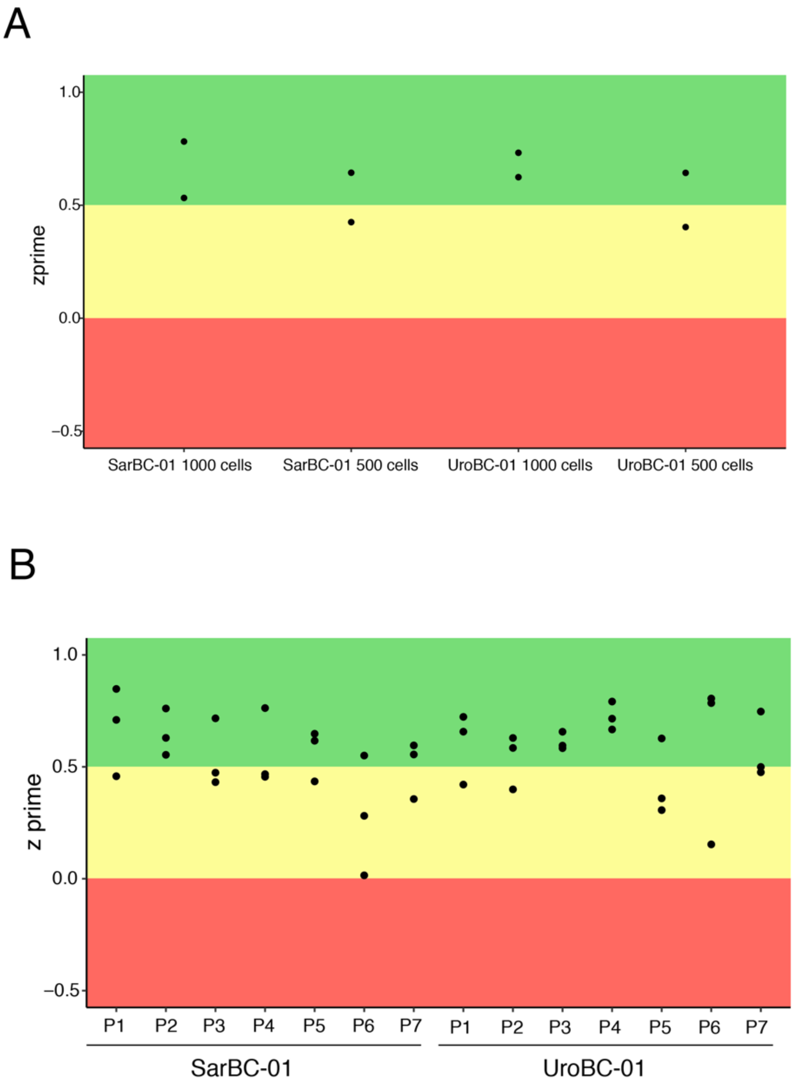


**Figure S5. Optimization and quality control of the high-throughput drug screen. (A)** For optimization of the screen, densities of 500 cells and 1000 cells per well were tested. Z prime values were calculated for each line for both densities. Assays with Z values above 0.5 are considered as excellent quality (green area), while assays with Z values between 0 and 0.5 are considered as good quality (yellow area).**(B)** For quality control of the screen, Z prime values were calculated based on 14 negative and positive controls for each plate (P1-P7, <0.01 % DMSO) for each line.

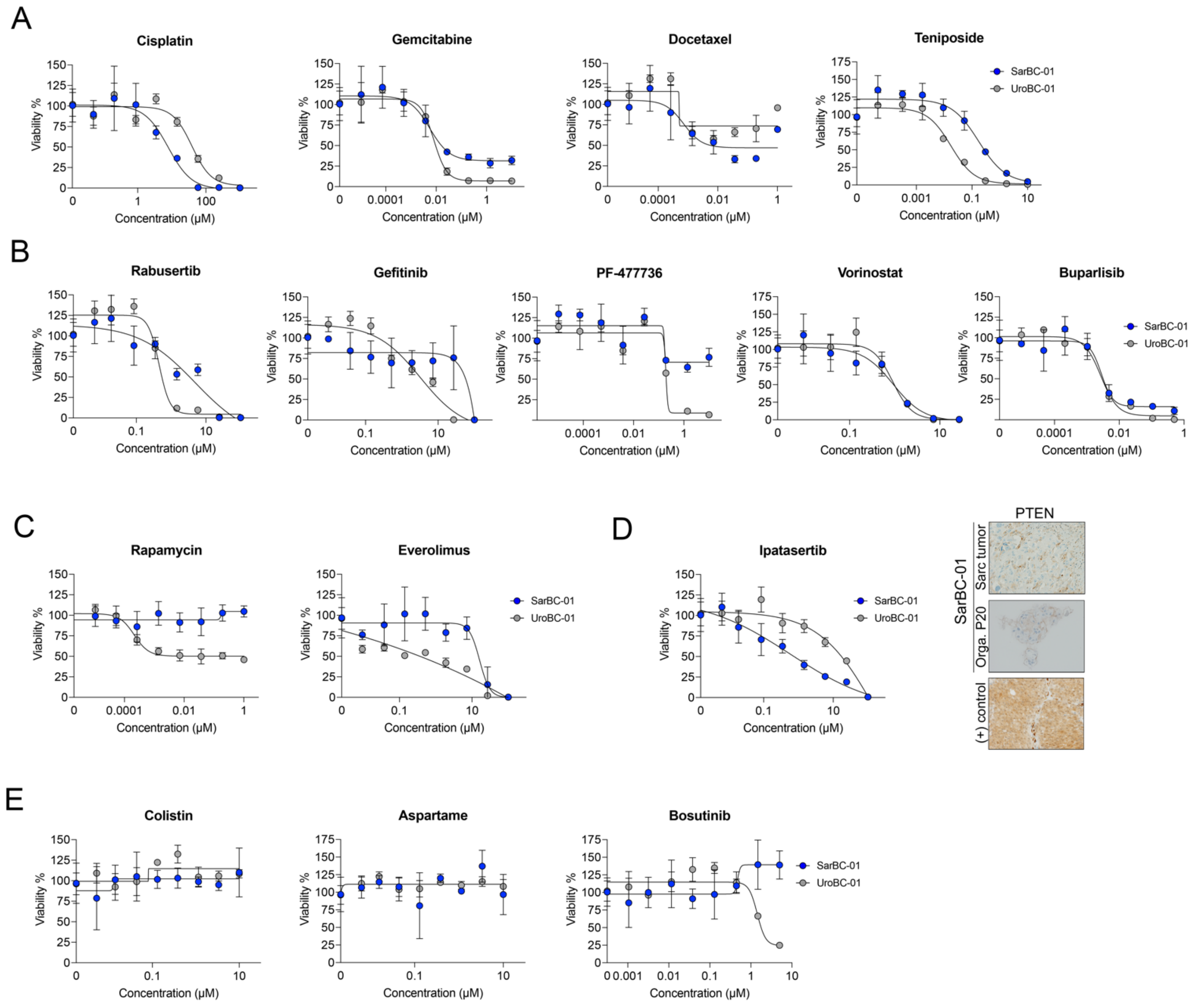


**Figure S6. Drug response analysis for selected compounds.** Dose responses curves for SarBC-01 (blue dots) and UroBC-01 (grey dots) organoids treated with a selected panel of drugs. Examples include **(A)** chemotherapeutic compounds, **(B)** targeted agents and **(C)** mTOR inhibitors. **(D)** SarBC-01 cells are responsive to the AKT inhibitor Ipatasertib (**left panel**), which may be explained by a loss of PTEN expression at protein level (**right panel**). Positive control for intact PTEN expression is also shown. **(E)** In the initial drug screen, a series of 17 specific compounds was identified as "ant-hits" as they seemingly enhanced the viability of SarBC-01 cells *in vitro* (**Fig. 3C**, group 8;). In dose-response analysis, no significant increase was observed for Colistin and Aspartame but SarBC-01 cells show slight increased viability at high concentrations of Bosutinib. $IC_{50}$ values for all tested compounds are reported in **Supplementary Table 3**, whenever available.

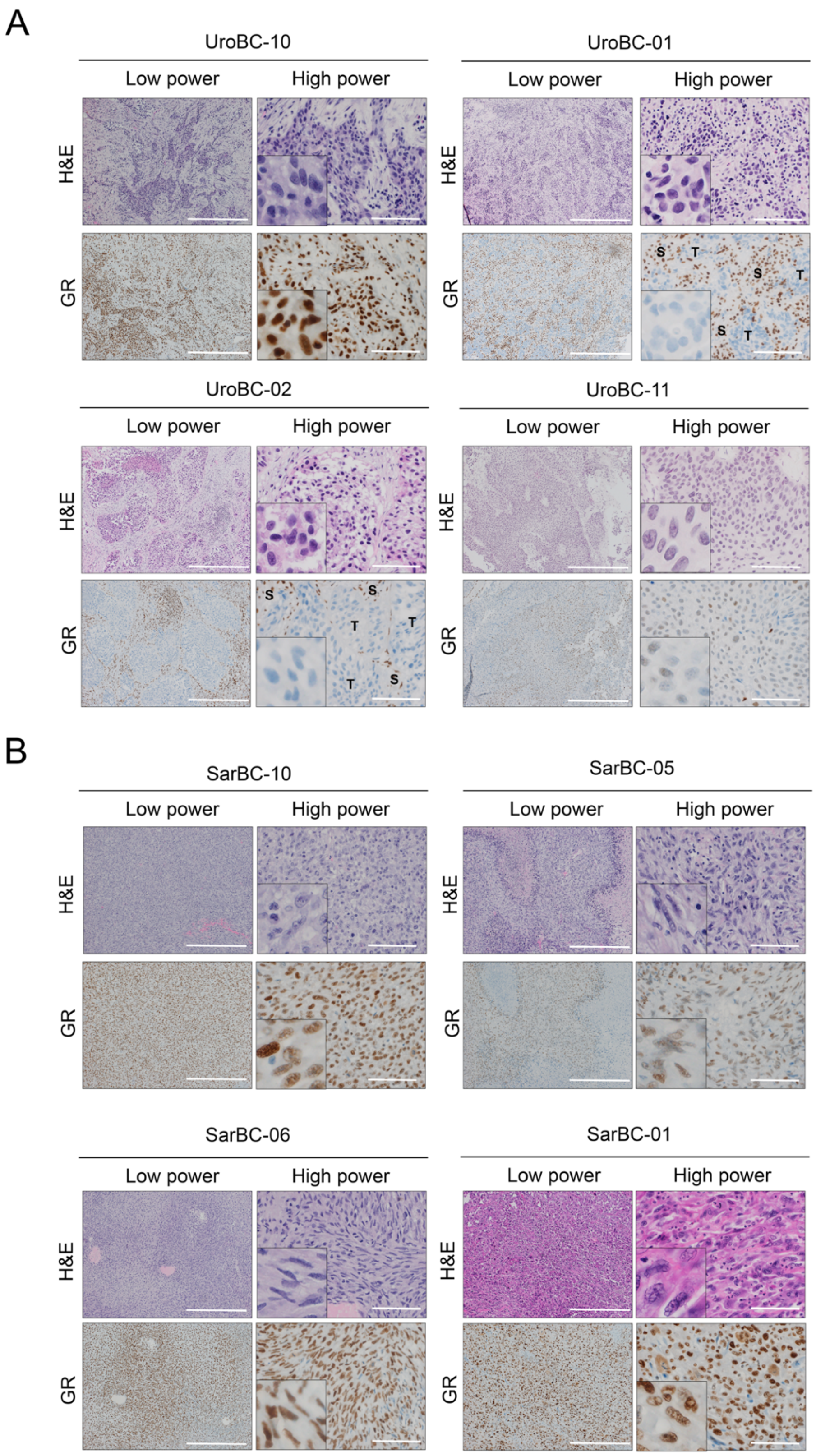


**Figure S7. Glucocorticoid Receptor (GR) expression in UroCa and SARC samples.** H&E and immunohistochemical staining for GR in UroCa samples **(A)** and SARC samples **(B).** *S: Stroma, T: tumor.* Scale bars represent 100 μm for high magnification images and 500 μm for low magnification images.

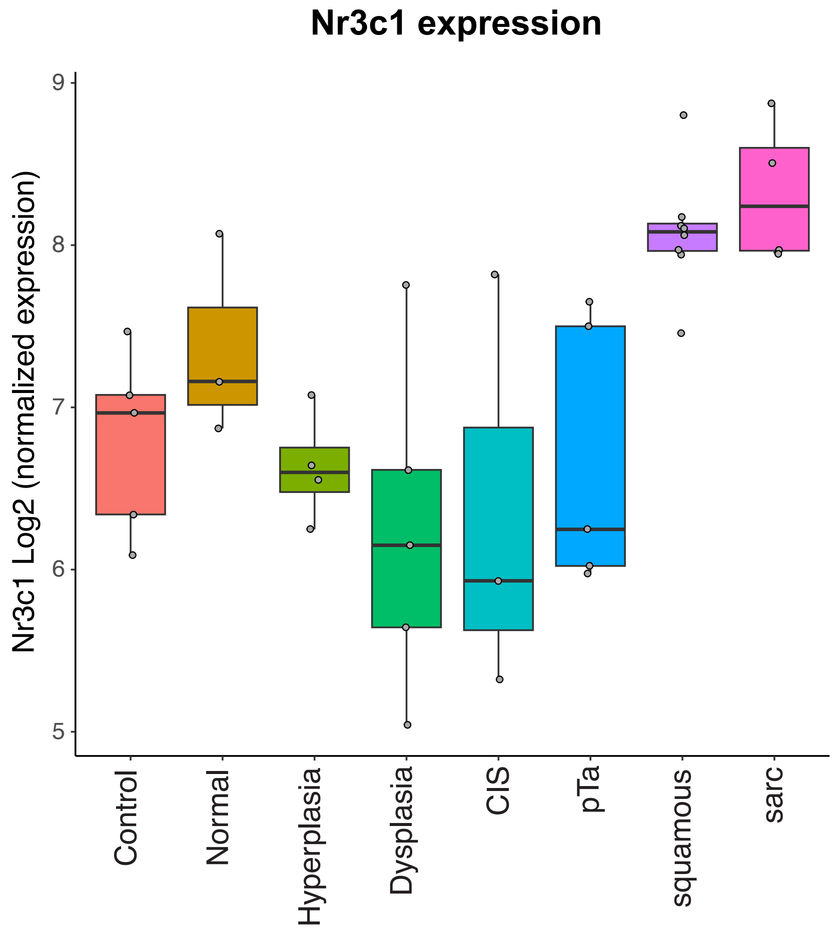


**Figure S8. Expression of *Nr3c1* in a mouse model of bladder cancer progression.** Data were obtained from a dataset downloaded from *Fontugne et al., 2023.* Shown are samples from mouse controls (i.e. not BBN treated, n=5), normal (BBN treated with normal histology, n=3), hyperplasia (n=4), dysplasia (n=4), carcinoma-in-situ (CIS, n=3), pTa (low-garde UroCa, n=5), basal/squamous (n=8), and sarcomatoid BC (sarc, n=4).

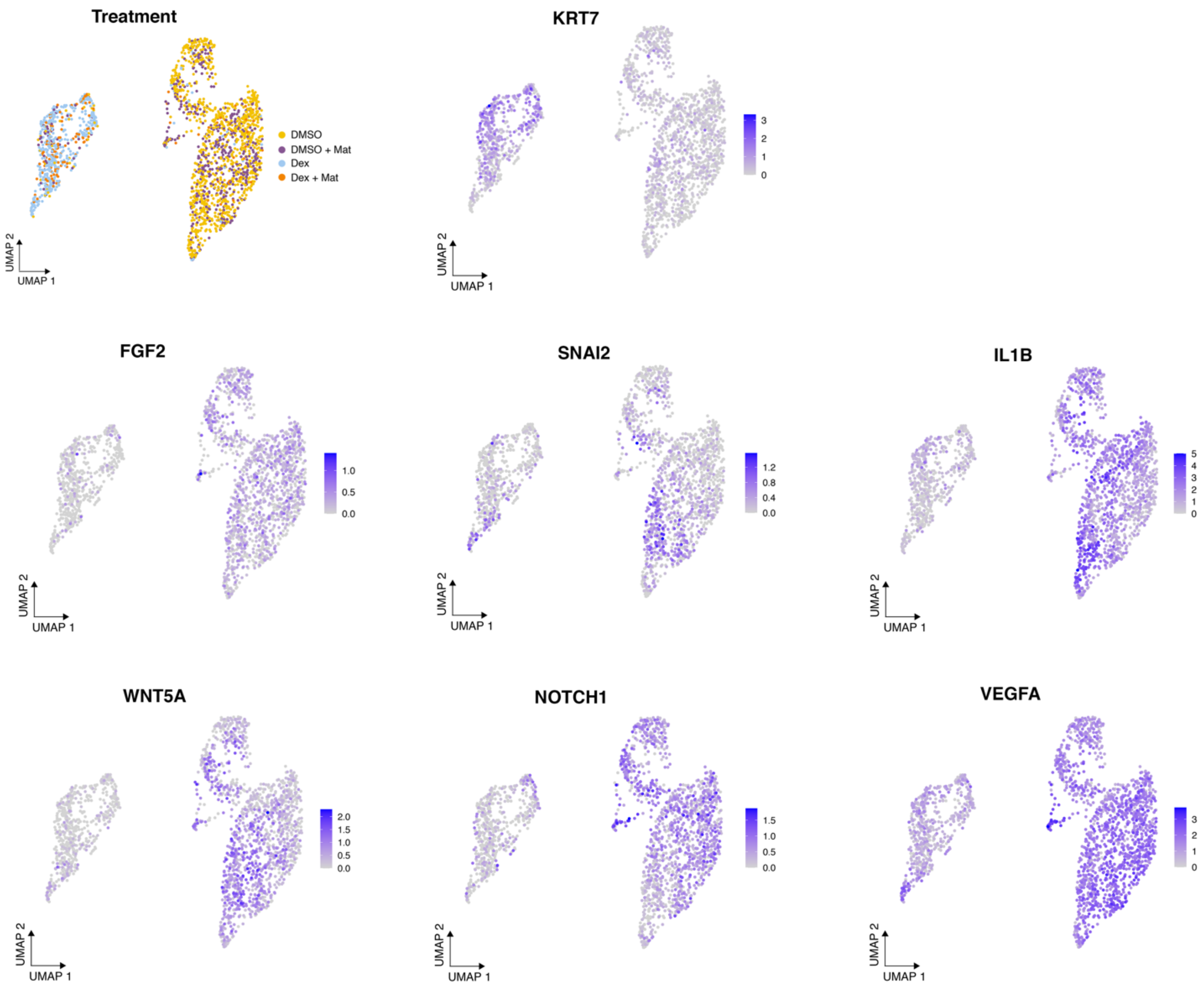


**Figure S9. Single-cell RNA sequencing of SarBC-01 cells following Dexamethasone treatment.** Shown are UMAP representations of the treatment conditions and expression of selected indicated genes. 2170 cells were analyzed following incubation with DMSO or Dexamethasone in invasion assay conditions and in presence of absence of Matrigel, leading to 4 different culture conditions (UMAP "treatment": DMSO, DMSO+Mat, Dex, Dex+Mat; see Material and Methods for more details).

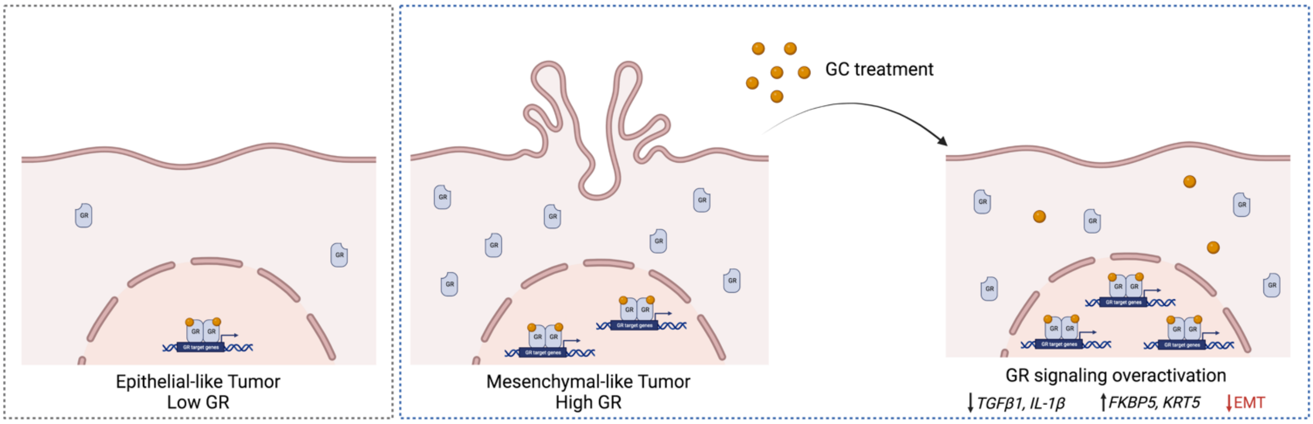


**Figure S10. Hypothetical model: Glucocorticoid Receptor signaling in SARC tumors.** Mesenchymal-like bladder tumors such as SARC, which display increased glucocorticoid receptor (GR) expression, may be more sensitive to glucocorticoid (GC) treatment. Treatment with GC leads to overactivation of the GR signaling pathway, as observed by *FKBP5* increase; this is accompanied by a reduction of expression of genes associated with EMT (e.g. *TGFB1*), an increase expression of epithelial-associated genes (e.g. *KRT5, KRT7*), and the acquisition of epithelial-like morphological features. *Created with BioRender.com.*
